# Lentiviral vector optimization enhances the expression and cytotoxicity of chimeric antigen receptors

**DOI:** 10.1101/2021.07.11.451925

**Authors:** Changjiang Guo, Han Chen, Jie Yu, Hui Lu, Xiali Guo, Xiaojuan Li, Tong Wang, Lingtong Zhi, Zhiyuan Niu, Wuling Zhu

## Abstract

Adoptive chimeric antigen receptor (CAR)-modified T or NK cells (CAR-T/NK) have emerged as a novel form of disease treatment. Lentiviral vectors (LVs) are commonly employed to engineer T/NK cells for the efficient expression of CARs. This study reported for the first time the influence of single-promoter and dual-promoter LVs on the CAR expression and cytotoxicity of engineered NK cells. Our results demonstrated that the selected CAR exhibits both a higher expression level and a higher coexpression concordance with the GFP reporter in HEK-293T or NK92 cells by utilizing the optimized single-promoter pCDHsp rather than the original dual-promoter pCDHdp. After puromycin selection, the pCDHsp produces robust CAR expression and enhanced *in vitro* cytotoxicity of engineered NK cells. Therefore, infection with a single-promoter pCDHsp lentivector is recommended to prepare CAR-engineered cells. This research will help to optimize the production of CAR-NK cells and improve their functional activity, to provide CAR-NK cell products with better and more uniform quality.

## Introduction

Therapies based on genetically modified immune cells are becoming effective agents in the fight against malignancies and other diseases ^1^. Chimeric antigen receptors (CARs) are functional artificial synthetic fusion proteins and designed to improve the targeting precision and efficacy of these therapies ^2^. A typical CAR is composed of an antigen recognition region, a transmembrane region, and several intracellular signaling domains. The antigen recognition domain is usually the single-chain variable fragment (scFv) or the ligand-binding domain of the natural receptor ^3^. CAR-modified T or NK cells can perform antigen recognition to specifically kill target cells such as cancer cells. The introduction of CAR-T/NK cell immunotherapy has become a milestone in the field of immunotherapy.

Viral vectors provide an efficient means for the modification of mammalian cells and are widely used in the preparation of CAR-T/NK cells ^4^. As one of the most extensively investigated viral vectors, lentiviral vectors (LVs), derived from the human immunodeficiency virus, are commonplace for both research and clinical applications. LVs can effectively transfer exogenous genes, such as CARs, into various cell types and lead to long-term gene expression due to stable integration into the genome ^5^. Previous studies have shown that the CAR-positive rate of CAR-T cells without enrichment prepared using LVs is mostly around 50% or even lower ^6–9^. For NK cells, which are notoriously difficult to transduce ^10^, the CAR-positive rate achieved with LVs tends to be lower ^11^. To overcome the defect of limited CAR expression in T cells, especially in NK cells, it is necessary to optimize the structure of the employed lentiviral vectors, in addition to improving the production and infection conditions of the lentivirus ^5,12^.

As a common series of lentiviral vectors lentiviral vector, the pCDH vectors are often harnessed for the transduction of both CAR-T ^13,14^ and CAR-NK cells ^15,16^. Each pCDH lentivector typically contains one or two constitutive promoters (e.g. CMV, EF1, and PGK) for stable expression of exogenous genes, fluorescent reporter genes, or resistance genes. For example, the dual promoter vector pCDH-CMV-MCS-EF1-copGFP-T2A-Puro is also a dual reporter construct and suitable for CAR-T/NK cell preparation. This vector is characterized by the presence of two independent promoters, the CMV promoter driving the gene of interest (GOI, e.g. CAR) cloned in the multiple cloning site (MCS) and the EF1 promoter driving the expression of the two markers (GFP and Puro). The GFP reporter gene indicates expression of the target gene by visible green fluorescence, while the Puro gene provides puromycin resistance for selection of stable clones. A self-cleaving T2A peptide sequence from the insect virus *Thosea asigna*, a molecular element used successfully to generate multiple proteins from a single promoter ^17^, is incorporated to mediate the co-expression of the GFP gene and the Puro gene.

However, for dual promoter vector pCDH-CMV-MCS-EF1-copGFP-T2A-Puro, the expression of the two markers is well coupled by the T2A peptide, but the expression of the gene of interest and the markers driven independently may differ depending on promoter activity, as described in the User Manual. What’s worse, we found that after infecting the NK92 cells with the vector pCDH-CMV-MCS-EF1-copGFP-T2A-Puro containing a CAR and screening them by puromycin selection, the enriched NK92 cells exhibited enhanced green fluorescence, while surface expression of the CAR did not increase as expected. This indicates that the dual promoter vectors are not suitable for the enrichment of engineered cells with high CAR expression by resistance screening.

Here, we further elucidated the effect of dual promoter lentiviral vectors on the preparation of CAR-engineered cells and constructed an optimized vector by replacing the second promoter with a self-cleaving P2A peptide. The results revealed intriguing differences in the expression of CAR and GFP in the engineered cells by the two different vectors with or without resistance screening. This study will help to construct optimized CAR-tailored lentiviral vectors for the preparation of CAR-engineered cells with high activity.

## Materials and Methods

### Cell lines and cell culture

The HEK-293T cell line, NK92 cell line, and human gastric carcinoma cell lines (SGC-7901 and KATO-III) were obtained from American Type Culture Collection (ATCC; Manassas, USA). NK92 cells were cultured in alpha modification of Eagle’s minimum essential medium (α-MEM; Hyclone, USA) supplemented with 12.5% horse serum (Solarbio, China), 12.5% fetal bovine serum (FBS; Biological Industries, Beit Haemek, Israel), 100 U/mL penicillin, 100 μg/mL streptomycin (Beyotime Biotechnology, China), and 100 U/mL recombinant human IL-2 (Peprotech, USA). HEK-293T cells were cultured in Dulbecco’s modified Eagle medium (DMEM; Hyclone, USA) supplemented with 10% FBS, 100 U/mL penicillin, and 100 μg/mL streptomycin. SGC-7901 cells were cultured in RPMI-1640 medium (Hyclone, USA) containing the same supplements as for DMEM. KATO-III cells were cultured in Iscove’s Modified Dulbecco’s Medium (IMDM; Hyclone, USA) containing the same supplements as for DMEM.

### pCDH lentivector optimization

The backbone of the original plasmid pCDH-CMV-MCS-EF1-copGFP-T2A-Puro (System Biosciences, Palo Alto, USA) was amplified and linearized via PCR using the high-fidelity PrimeSTAR GXL DNA Polymerase (Takara Bio, Japan). In the PCR product, the EF1 promoter was removed and the self-cleaving peptide P2A sequence was introduced. The recombinant plasmid was obtained by ligation using Seamless Cloning Kit (Beyotime Biotechnology, China) with 37-bp overlap sequences flanking the amplification product and then verified by sequencing. The optimized plasmid vector was referred to as pCDH-CMV-MCS-P2A-copGFP-T2A-Puro. Primer sequences are as follows:

5’-CAGCCTGCTGAAGCAGGCTGGAGACGTGGAGGAGAACCCTGGACCT

ATGGAGAGCGACGAGAGCGG-3’ (forward primer)

5’-CAGCCTGCTTCAGCAGGCTGAAGTTAGTAGCTCCGCTTCCGCTAGC

GCGGCCGCGGATCCGATTTA-3’ (reverse primer)

### NKG2D-based CAR construction

The synthetic chimeric antigen receptor (CAR) was constructed by fusing the extracellular domain of human NKG2D to the IgG4 hinge region, the transmembrane domain of human CD28, and the cytoplasmic domains of human CD28, CD137, and CD3ζ. This CAR construct was subcloned into the original and optimized lentiviral plasmid vectors with standard molecular biology techniques. The CAR sequence was preceded by the Flag tag for detection.

### Transient transfection of HEK-293T cells

All lentivector constructs were transfected into HEK-293T cells using Calcium Phosphate Cell Transfection Kit (Beyotime Biotechnology, China) according to manufacturer instructions. Analysis of transfected cells was performed at 48 h post-transfection.

### Lentivirus Production

Each lentivector plasmid was co-transfected into HEK-293T cells with packaging plasmid psPAX2 and pMD2.G at 4:3:1 ratio (1000ng:750ng:250ng, respectively) Calcium Phosphate Cell Transfection Kit. Lentivirus particles were harvested from the supernatant at 48 h and 72 h post-transfection and concentrated by incubation with PEG 8,000 (80 g/L) and NaCl (17.5 g/L) during overnight shacking (100 rpm) at 4°C, followed by centrifugation for 30min at 3500×g. The pellet was then resuspended in 1 mL phosphate buffer saline (PBS) and stored at −80°C until further processing. Lentivirus titers were measured by using the qPCR Lentivirus Titer Kit (Applied Biological Materials, China).

### Production of CAR-NK92 cells

NK92 cells were transduced with each pCDH lentivirus (multiplicity of infection is 30) in the presence of 8 μg/mL polybrene (Sigma). Then, the cells continue to expand at 37°C. The expression of CAR and GFP was determined 72 hours after transduction. The transformants were selected for two weeks with 800 ng/mL puromycin (Solarbio, China).

### Flow cytometry

Analysis of CAR expression on transduced NK92 cells was done by staining with APC-conjugated anti-human CD314 (NKG2D) mAbs and anti-Flag Tag mAbs (Biolegend, USA). NKG2DLs expression in each cancer cell line was stained with recombinant human NKG2D Fc chimera (R&D Systems, USA) and then FITC-conjugated anti-human Fc secondary antibody was used as a detection antibody (BD Biosciences, USA). Samples were analyzed by Guava^®^ easyCyte^™^ Flow Cytometer (Millipore, USA). Flow cytometry data were analyzed by FlowJo V10 software (FlowJo LLC, USA).

### Cytotoxicity assays

Cytotoxicity of transduced NK92 cells was determined by an LDH release assay. LDH release assays were done using the LDH Cytotoxicity Assay Kit (Beyotime Biotechnology, Shanghai, China) according to manufacturer instructions. The NK92 cells and tumor cells were co-cultured in the α-MEM medium at the effector-to-target ratio (E:T ratio) of 5:1 in triplicates in 96-well flat-bottom plates for 4 hours. The percentage of specific lysis was calculated according to the following formula: Cytotoxicity (%) = (experimental LDH release - spontaneous LDH release)/(maximal LDH release - spontaneous LDH release) × 100.

### Cytokine assays

The NK92 cells were co-cultured with tumor cells in α-MEM complete medium at the E:T ratio of 5:1 in triplicate wells. Culture supernatant was harvested after 12 hours, and the concentrations of IFN-γ, TNF-α, Granzyme B and perforin were determined by ELISA kits (ABclonal, Biolegend, and Abcam) according to the manufacturer’s instructions.

### Statistical analysis

All in vitro experiments were performed at least in triplicate. Data were presented as mean ± standard deviation (SD). The differences between groups using Student’s t-test or Wilcoxon rank-sum test, as indicated in each figure legend. By convention, we referred to statistically significant as P < 0.05 and statistically highly significant as P < 0.001. Statistical analysis was performed with GraphPad Prism software version 6.0 (GraphPad Software Inc., La Jolla, USA).

## Results

### Optimization of the dual promoter lentivector pCDH-CMV-MCS-EF1-copGFP-T2A-Puro and construction of an NKG2D-based CAR

To improve the co-expression concordance of the gene of interest with the reporters, we replaced the EF1 promoter in the dual promoter lentivector pCDH-CMV-MCS-EF1-copGFP-T2A-Puro (Fig. 1A) with the self-cleaving P2A peptide sequence and thus constructed a novel single promoter lentivector pCDH-CMV-MCS-P2A-copGFP-T2A-Puro (Fig. 1B). The unmodified and modified vectors were abbreviated as pCDHdp and pCDHsp, respectively. The pCDHsp links the gene of interest and the two reporter genes with two 2A peptides in a single transcript, so their expression is driven by the same CMV promoter upstream. Moreover, compared with the original pCDHdp, the pCDHsp was shortened by nearly 500 bp, thus can hold the insert with a larger size than the original pCDHdp (Fig. 1A-B and Fig. S1).

**Fig. 1.**
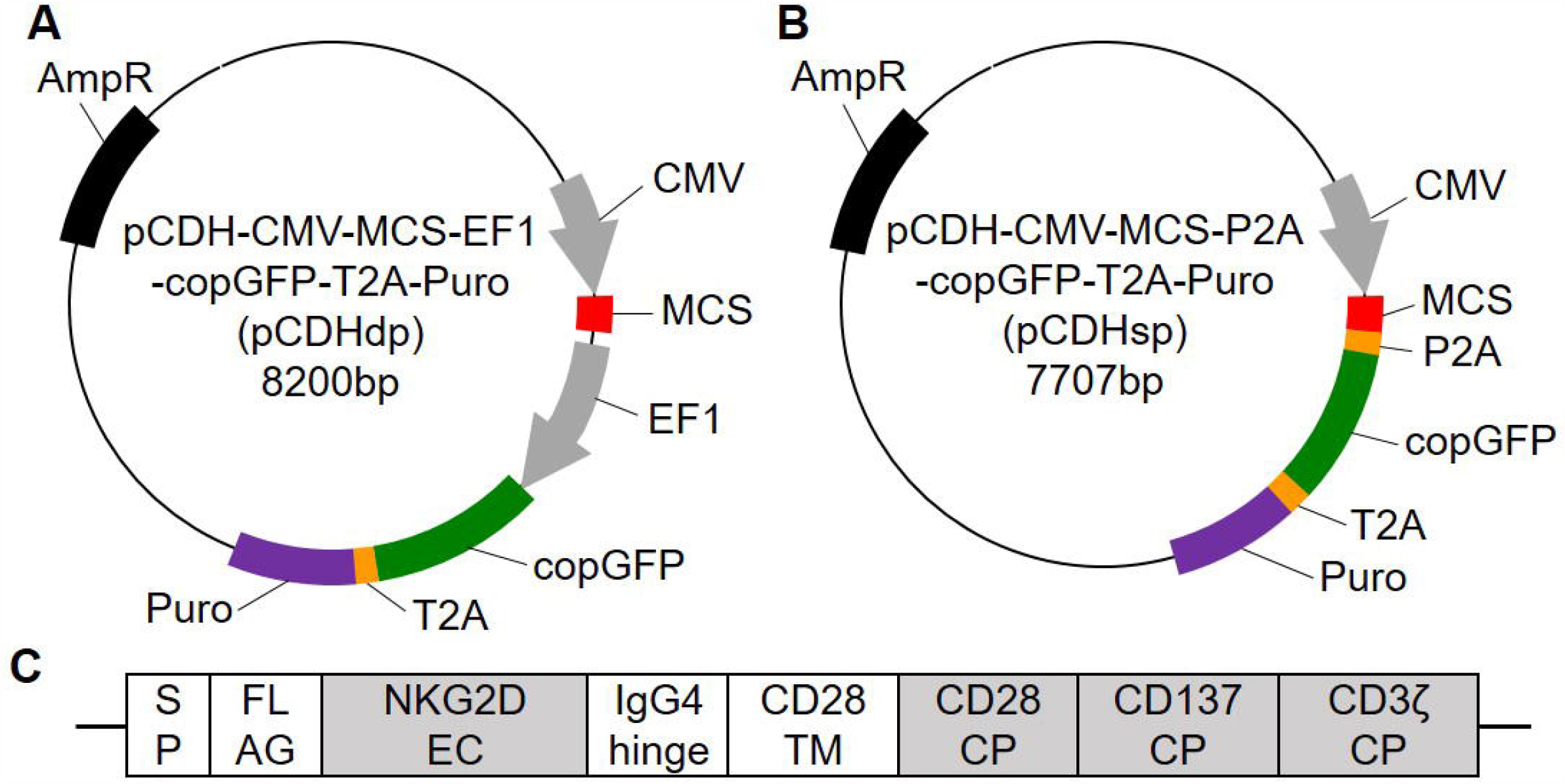
Schematic representation of original pCDH lentivector (A), optimized pCDH lentivector (B), and NKG2D-based chimeric antigen receptor (C). Multiple cloning site (MCS), signal peptide (SP), Flag tag (FLAG), extracellular (EC) domain, transmembrane (TM) domain, and cytoplasmic (CP) domains are indicated.

To clarify the differences in expression and function of the inserted gene between pCDHdp and pCDHsp, we next constructed an NKG2D-based CAR as the insert (Fig. 1C). NKG2D is a vital activating receptor in NK cells and its ligands (NKG2DLs) are up-regulated on stressed cells, which include tumor and infected cells ^18^. We incorporated this NKG2D-CAR into two pCDH lentivectors and generated recombinant vectors pCDHdp-CAR and pCDHsp-CAR, respectively.

### Expression of CAR and GFP in transfected HEK-293T cells

To confirm the expression of NKG2D-CAR and GFP, four plasmids (pCDHdp, pCDHsp, pCDHdp-CAR, and pCDHsp-CAR) were transfected into the HEK-293T cell line, respectively. All of the HEK-293T cells emitted the green fluorescence at 48 hours post-transfection under the fluorescence microscopy, the weakest fluorescence intensity was observed in pCDHsp-CAR transfected HEK-293T cells (Fig. 2A). Consistently, flow cytometric analysis showed that HEK-293T cells transfected with pCDHsp-CAR had the lowest GFP-positive expression rate (Fig. 2B) and the lowest mean fluorescence intensity (Fig. 2C).

**Fig. 2.**
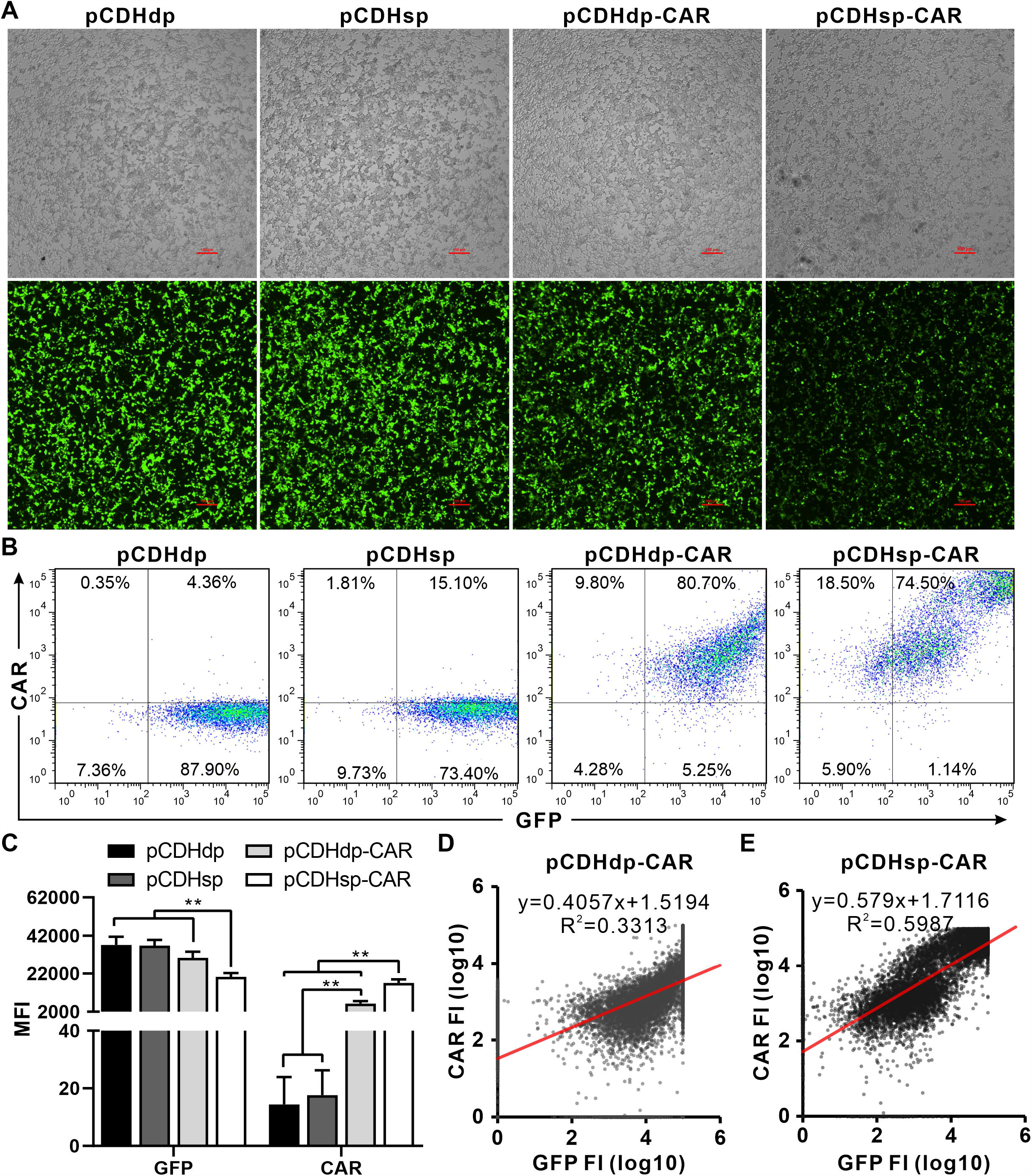
Expression of GFP and CAR in HEK-293T cells transfected with four pCDH lentiviral vectors. (A) Microscopical images of transfected HEK-293T cells in the bright field channel (top row) and GFP channel (bottom row) at 48 hours post-transfection. Scale bar: 100 μm. (B-C) Expression of GFP and surface expression of CAR in transfected in HEK-293T cells, as assessed by flow cytometry. Each MFI was determined by subtraction of the unstained MFI value. MFI, mean fluorescence intensity. Data are shown as mean ± SD. Statistical analysis was done by two-tailed Student’s t-test; **p < 0.01. (D-E) Relationship between the CAR expression level and the GFP fluorescence level on a log_10_-log_10_ scale in the HEK-293T cells transfected by two CAR-carrying pCDH vectors. FI, fluorescence intensity.

Meanwhile, HEK-293T cells transfected with the two CAR-carrying pCDH vectors had the significant surface expression of CAR compared with empty vectors(Fig. 2C), and the positive expression rate of both was above 90% (Fig. 2B). Importantly, the mean fluorescence intensity (MFI) showed that the expression level of CAR in pCDHsp-CAR transduced HEK-293T cells was significantly higher than that in pCDHdp-CAR (p = 0.00126) and empty vectors (p < 0.0001) (Fig. 2C). These results indicated that replacing the EF1 promoter with the P2A peptide significantly enhanced CAR expression, but conversely impairs GFP expression (Fig. S2A). We then analyzed the co-expression concordance between GFP fluorescence and CAR by linear regression. As expected, there was a higher expression correlation between GFP and CAR in pCDHsp-CAR transduced HEK-293T cells than that in pCDHdp-CAR (R^2^: 0.60 vs 0.33; Fig. 2D-E, Fig. S2B-C). Thus, the linking of the P2A peptide conferred better-coupled expression between CAR and the downstream GFP reporter in the optimized pCDHsp vector.

### Expression of GFP and CAR in stably transfected NK92 cells

Lentiviral particles generated from four plasmids (pCDHdp, pCDHsp, pCDHdp-CAR, and pCDHsp-CAR) were harnessed to transduce NK92 cells separately, and then puromycin selection was performed to achieve stable transfection of NK92 cells. We monitored the expression of GFP and CAR by flow cytometry during puromycin screening. The results showed that GFP fluorescence in transfected NK92 cells was enhanced as the screening proceeded (Fig. 3A-B, Fig. S3). After 14 days of screening, the green fluorescence in the two empty vector-transfected NK92 cells was significantly higher than that in the transfected NK92 cells of the two CAR-carrying vectors (p < 0.001; Fig. 3B). This indicated that insertion of the CAR into the pCDH vector in NK92 cells interferes with the expression of the GFP reporter. Interestingly, the MFI data showed that while the green fluorescence was weakest in NK92 cells stably transfected with pCDHsp-CAR, their fluorescence had the highest increase by 9.3-fold compared to unscreened cells (Fig. 3B). These results indicated that the insertion of CAR into the pCDHdp or pCDHsp significantly reduces GFP fluorescence.

**Fig. 3.**
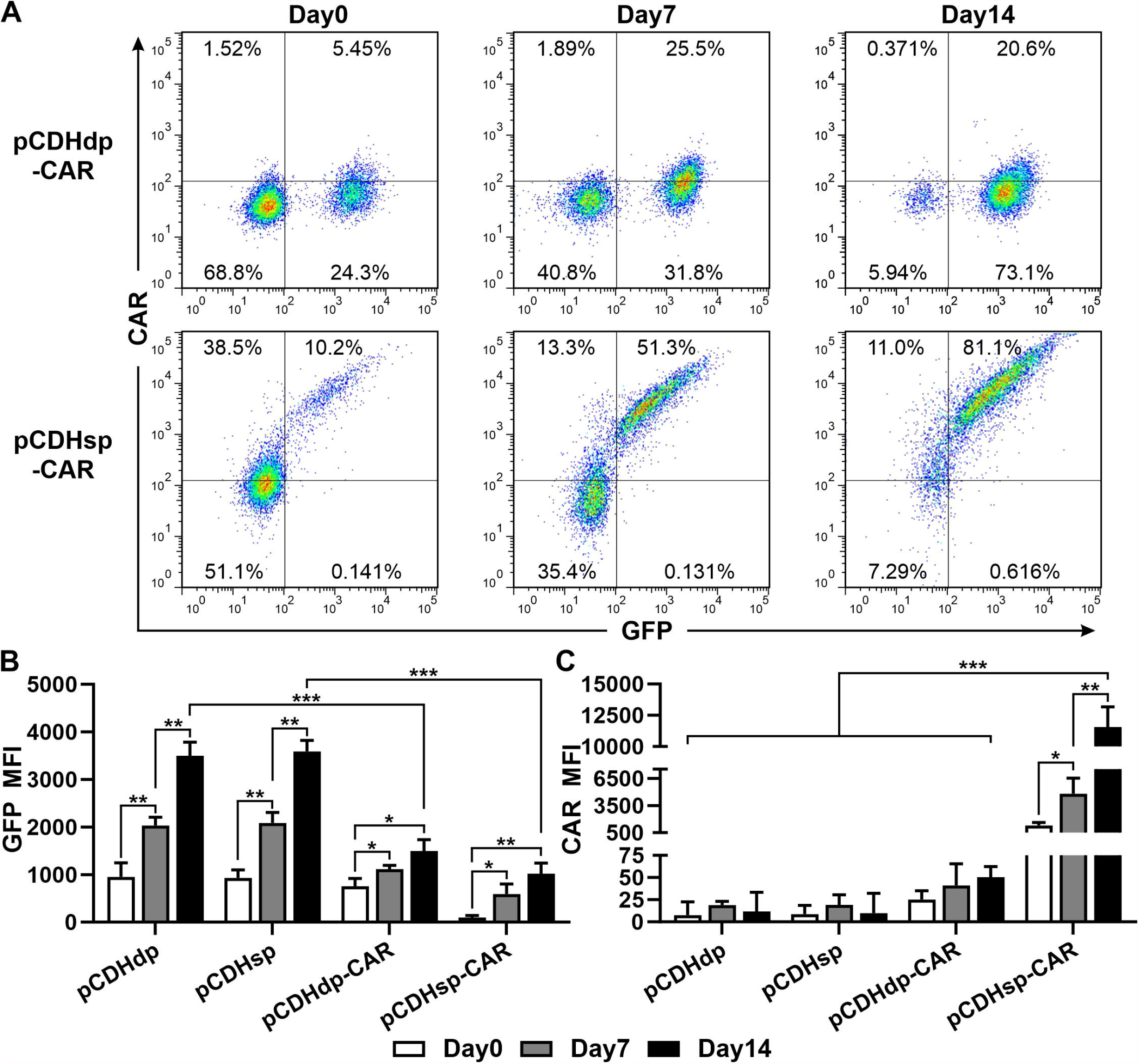
Expression of GFP and surface expression of CAR in NK92 cells transfected with four pCDH lentiviral vectors, as assessed by flow cytometry. (A) Flow cytometric analysis of GFP and CAR in the NK92 cells transfected by pCDHdp-CAR and pCDHsp-CAR during puromycin screening. (B-C) The mean fluorescence intensity (MFI) of GFP (B) and CAR (C) in transfected NK92 cells during puromycin screening. Each MFI was determined by subtraction of the unstained MFI value. Day0, the day before the start of puromycin selection. Day7 / Day14, day 7 and 14 after the start of puromycin selection. MFI, mean fluorescence intensity. Data are shown as mean ± SD. Statistical analysis was done by two-tailed Student’s t-test; *p < 0.05, **p < 0.01, ***p < 0.001.

Meanwhile, CAR expression in transfected NK92 cells was evaluated by flow cytometry. The assay data showed that CAR expression (both positivity rate and expression intensity) in pCDHsp-CAR transfected NK92 cells increased continuously with the puromycin screening (Fig. 3A&D). After 7 and 14 days of screening, the expression of CAR increased to 2.72-fold and 7.97-fold, respectively. Whereas, CAR expression in pCDHdp-CAR transfected NK92 cells exhibited only a weak increase in positivity rate early in the screening (Fig. 3B&D), while CAR expression in NK92 cells transfected with the empty vectors pCDHdp and pCDHsp did not change significantly (Fig. 3C, Fig. S3). Following 14 days of puromycin selection, the pCDHsp-CAR transfected NK92 cells emitted the weakest green fluorescence as reflected by the MFI data (Fig. 3B) and fluorescence microscopy observations (Fig. 4A). Nevertheless, pCDHsp-CAR transfected NK92 cells had a CAR-positive rate of 92.1%, as compared to 21.0% for pCDHdp-CAR (Fig. 3A). Crucially, the surface expression level of CAR in pCDHsp-CAR transfected NK92 cells was 51.26-fold higher than that in pCDHdp-CAR transfected NK92 cells before screening and increased to 229.41-fold after 14 days of screening (Fig. 3D). These results suggested that the optimized pCDHsp vector can help tremendously to improve the expression of CAR in NK92 cells before and after puromycin selection at the cost of impairing the expression of GFP (Fig. S4).

**Fig. 4.**
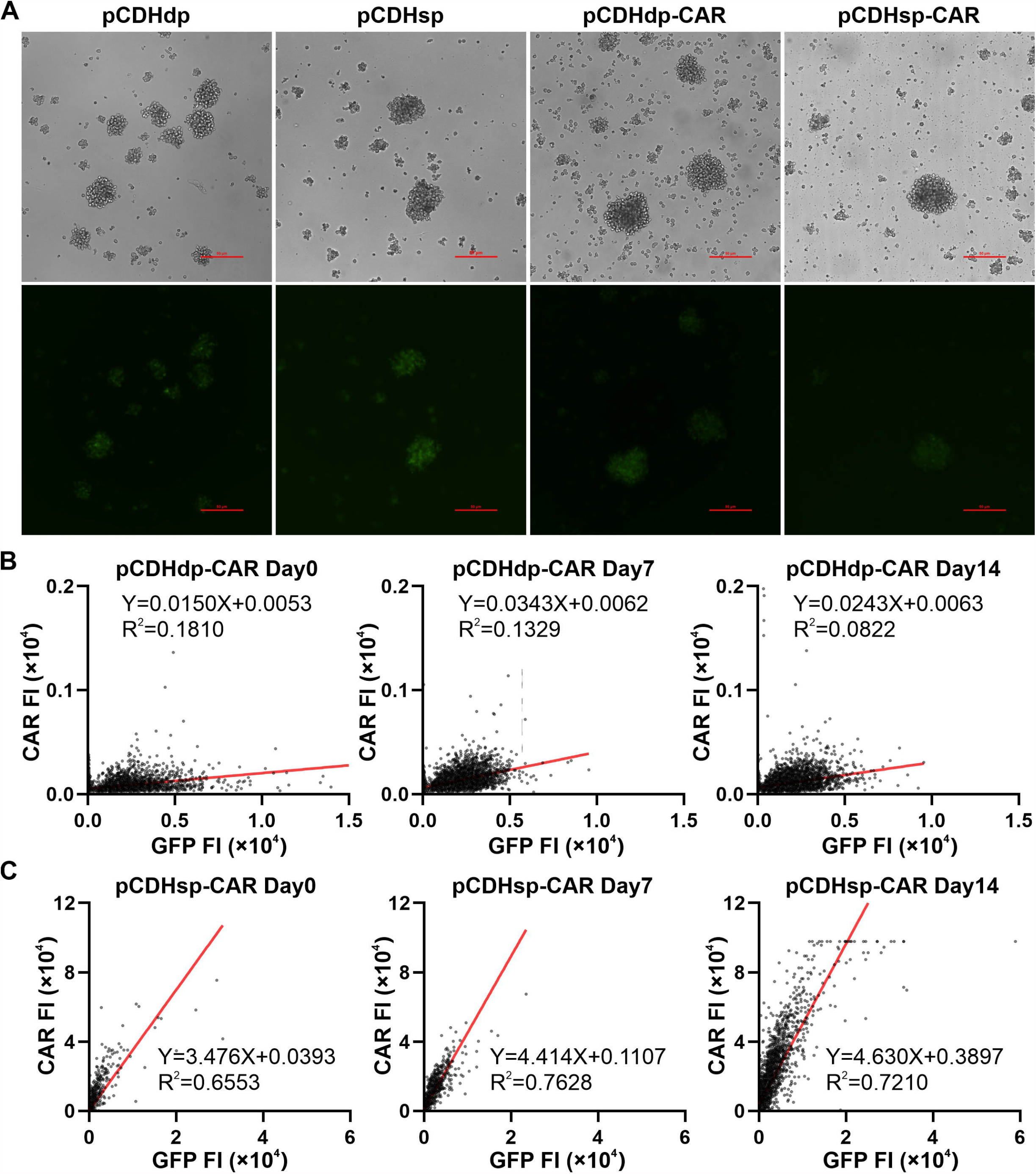
Green fluorescence and co-expression of GFP and CAR in transfected NK92 cells. (A) Microscopical images of stably transfected NK92 cells in the bright field channel (top row) and GFP channel (bottom row). Scale bar: 50 μm. (B-C) Relationship between the CAR expression level and the GFP fluorescence level in the NK92 cells transfected by two CAR-carrying pCDH vectors during puromycin selection. FI, fluorescence intensity. Day0, the day before the start of puromycin selection. Day7 / Day14, day 7 and 14 after the start of puromycin selection.

We then analyzed the co-expression concordance between GFP and CAR using linear regression. There was no significant co-expression of GFP and CAR in pCDHdp-CAR transfected NK92 cells with or without puromycin selection (R^2^ < 0.2; Fig. 4B, Fig. S5A). In contrast, for pCDHsp-CAR transfected NK92 cells, there was a significant co-expression concordance between GFP and CAR (R^2^ > 0.65; Fig. 4C, Fig. S5B). Meanwhile, the increase of both slope and intercept of the regression line following puromycin selection indicated that the expression of CAR was improved at a higher rate than that of GFP (Fig. 4C). Consequently, the optimization of dual-promoter pCDH lentivector conferred a highly concordant coexpression of CAR and the reporters, which is very helpful for precise detection of CAR expression by green fluorescence and for enriching cells with robust CAR expression by resistance selection.

### In vitro cytotoxicity of NK92 cells stably expressing CAR

To investigate the cytotoxicity of NK92 cells stably expressing NKG2D-CAR against tumor cells, two human gastric carcinoma cell lines KATO-III and SGC-7901 with different surface expression levels of NKG2D ligands (NKG2DLs) were used as killing targets. The flow cytometry results showed a very limited expression of NKG2DLs in KATO-III cells and a significantly higher expression of NKG2DLs in SGC-7901 cells (8.03% vs 57.6%; Fig. 5A). We then investigated the cytotoxicity of untransduced and stably transduced NK92 cells against these cancer cells by LDH release assay *in vitro*. The results demonstrated that NKG2D-CAR significantly enhanced the cytotoxicity of NK92 cells on SGC-7901 cells with higher expression of NKG2DLs, but did not effectively augment the killing of KAOT cells (Fig. 5B). These results suggested that the effective killing of target cells by transfected NK92 cells is dependent on the binding of NKG2D to its ligands.

**Fig. 5.**
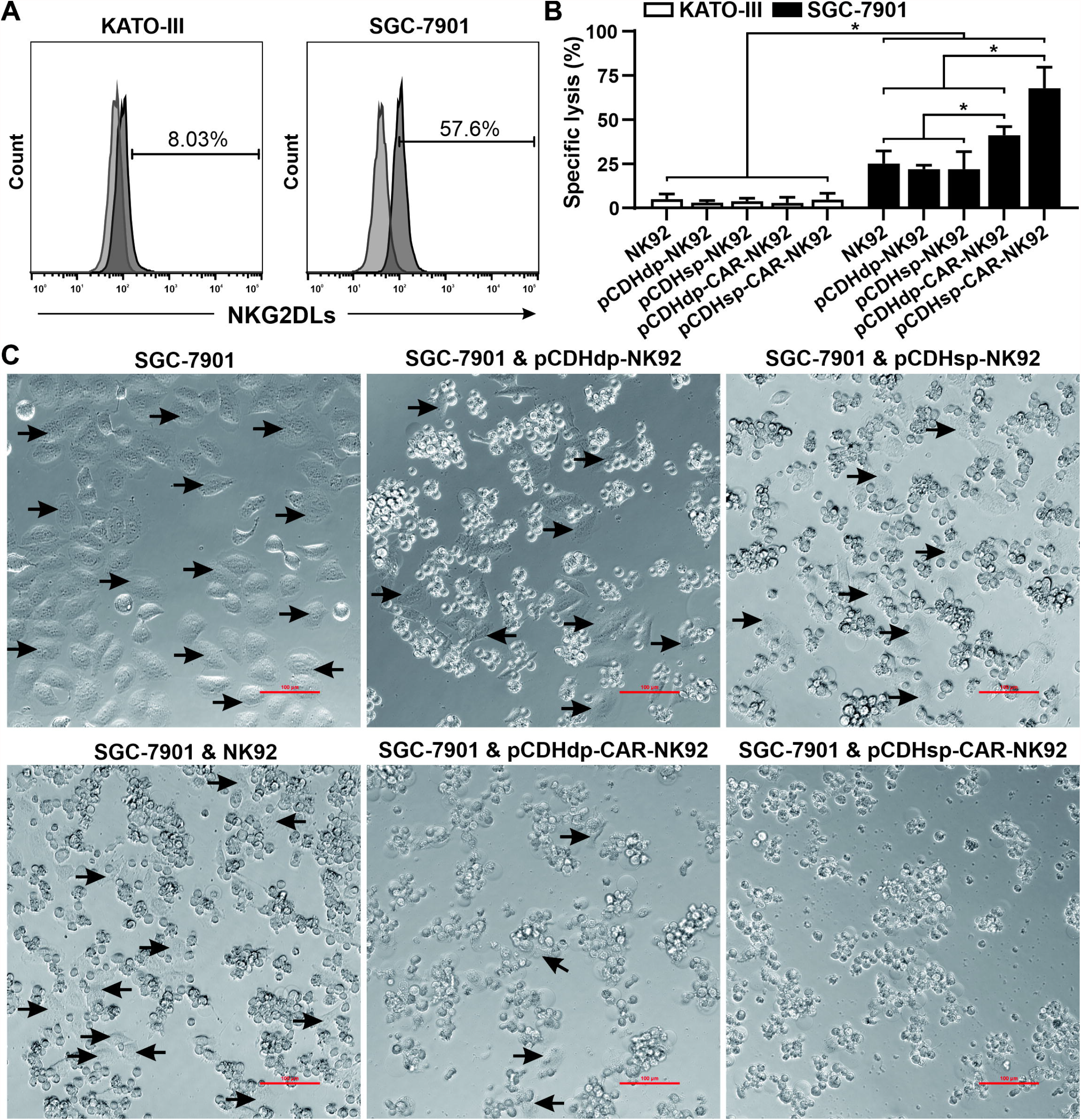
CAR-mediated cytotoxic activity in stable transfected NK92 cells. (A) Surface expression of NKG2D ligands (NKG2DLs) in the human gastric carcinoma cell lines KATO-III and SGC-7901, as assessed by flow cytometry. (B) LDH release cytotoxicity assays were performed to measure the specific lysis of KATO-III and SGC-7901 cells by untransduced or stably transduced NK92 cells at an effector to target ratio (E:T) of 5:1 after co-incubation for 4 hours. The mean percentages of specific cell lysis ± SD are shown. Statistical analysis was done by two-tailed Student’s t-test; *p < 0.05. (C) Microscopic observation of kinetic cytotoxicity of untransduced and transduced NK92 cells against SGC-7901 cells at the E:T ratio of 5:1. The wall-attached cancer cells are indicated by black arrows. Scale bar, 100 μm.

Meanwhile, limited expression of CAR in pCDHdp-CAR-NK92 cells was sufficient to present effective killing of SGC-7901 cells, which was further significantly enhanced in pCDHsp-CAR transfected NK92 cells (Fig. 5B-C). This indicated that more surface expression of CAR will enhance *in vitro* cytotoxicity of the modified cells. Furthermore, compared to the other transfected NK92 cells, there were almost no wall-attached SGC-7901 cells after 4 hours of co-incubation with pCDHsp-CAR-NK92 cells (Fig. 5C). Correspondingly, ELISA assays revealed increased secretion of multiple cytokines including IFN-γ, TNF-α, granzyme B, and perforin upon lysis of SGC-7901 cells by pCDHsp-CAR-NK92 cells (Fig. S6). These results indicated that the optimized pCDHsp vector not only facilitates the enhancement of CAR expression and co-expression with GFP but also significantly augments the function of the modified NK cells.

## Discussion

In this study, a dual promoter lentivector (pCDH-CMV-MCS-EF1-copGFP-T2A-Puro, pCDHdp) was optimized into a single promoter lentivector (pCDH-CMV-MCS-P2A-copGFP-T2A-Puro, pCDHsp) to provide a better gene transfer tool for CAR engineering. The pCDHsp with the self-cleaving P2A peptide instead of the EF1 promoter not only facilitated the enhancement of CAR expression but also significantly enhanced the co-expression of CAR and GFP in stably transfected NK92 cells derived from resistance selection. More importantly, the *in vitro* cytotoxicity of the CAR-engineered NK cells was thus significantly enhanced.

In recent years, CAR-engineered immune cell (e.g. T cells, NK cells) therapy has been gradually applied to treat cancers, autoimmune disorders, and viral infections, including SARS-CoV-2 ^19^. There are various gene delivery tools for CAR-T/NK cell preparation, including viral-based vectors (e.g. retroviruses, adenoviruses, lentiviruses) or nonviral vectors ^20^. As widely used gene transfer vectors for the production of CAR-T/NK cells, lentiviruses have a broad range of host cell tropism for infection and enable persistent and stable expression of the delivered genes. The high infection efficiency of produced CAR-T/NK cells is a key element of quality. Both the proportion and level of CAR surface expression in infected cells are more critical. However, the global CAR-positive rate in most of the infected cells was only about 50% due to the limitation of transfection efficiency and vector construct. Therefore, it is necessary to use a drug selection to increase the surface expression ratio and intensity of CAR in the infected cells.

Our study showed that when expressed with the reporter genes driven by separate promoters, CAR expression remained low in the stably transfected NK cells by puromycin selection (Fig. 3A&C). Meanwhile, we found that the expression of GFP, which is linked to the resistance gene by the T2A peptide, was significantly enhanced under the drive of the same EF1 promoter (Fig. 3A-B). This is due to the theoretical equimolar expression levels of 2A peptide-linked proteins, which are translated as a single unit. Thus, 2A peptides are commonly used for robust equimolar co-expression of two proteins, whereas the internal ribosome entry site (IRES) sequence allows the expression of two proteins at a ratio of approximately 3:1 ^21^. For these reasons, we replaced the EF1 promoter in the original pCDH lentivector with the P2A sequence, which is a self-cleaving peptide derived from *Porcine Teschovirus-1* and has the highest cleavage efficiency in human cell lines ^17^.

As expected, the P2A peptide was able to correlate GFP fluorescence with CAR expression (Fig. 2B&3A). More importantly, stably transfected NK cells by puromycin selection exhibited enhanced expression of both CAR and GFP. The robust expression of CAR augments NK cell-mediated cytotoxicity (Fig. 5B-C). Therefore, optimization of the dual promoter pCDH lentivector provides an effective method for monitoring CAR expression by green fluorescence and enables the enrichment of cells with robust CAR expression and increased cytotoxicity.

In conclusion, we modified a dual promoter lentivector into a single promoter one by the use of a P2A peptide. The optimized lentivector eliminated the difference in expression between the inserted CAR and the markers and achieved stable transduction of NK92 cells with robust expression of CAR and enhanced cytotoxicity. This research may provide a potential strategy to increase the therapeutic efficacy of the CAR-T/NK cells and efficiently enable the translation of CAR-based immunotherapy to the clinic.

## Authors contributions

C. Guo: designing experiments, data collection, analyzing and interpreting results, funding acquisition, writing the manuscript

H. Chen: carring out experiments, data collection, analyzing and interpreting results

J. Yu: data collection

H. Lu: data collection

X. Guo: data collection

X. Li: data collection

T. Wang: data collection

L. Zhi: supervising the project

Z. Niu: supervising the project

W. Zhu: supervising the project, funding acquisition, writing original draft

## Conflict of interest

The authors declare that there is no conflict of interest.

## Funding

This study was supported by the Major Science and Technology Projects in Henan Province (Grant No. 201100311100); the National Natural Science Foundation of China (Grant No. 82003261); the National Key R&D Program of China (Grant No. 2019YFA0906000); the Science and Technology Projects in Xinxiang (Grant No. 20GG006); the Key Scientific Research Project of Colleges and Universities in Henan Province (Grant No. 19A180026).

## Acknowledgments

The authors thank the Stem Cell and Biotherapy Engineering Research Center of Henan at Xinxiang Medical University for providing the lentivirus vector system.

## Abbreviations

CAR: chimeric antigen receptor
LV: Lentiviral vector
MFI: mean fluorescence intensity

## Notes

### Competing Interest Statement

The authors have declared no competing interest.

