## supplemental figures for "Lentiviral vector optimization enhances the expression and cytotoxicity of chimeric antigen receptors"

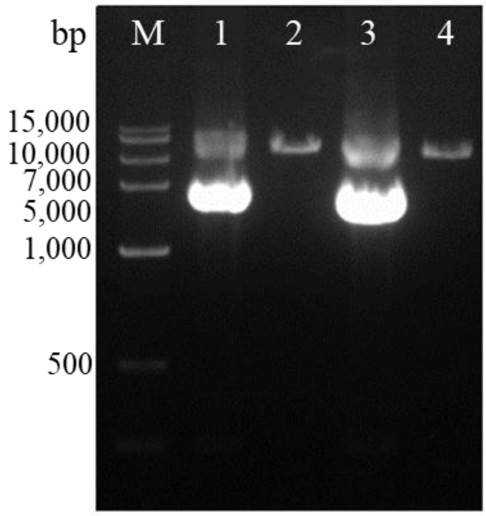


**Fig. S1**. Agarose gel electrophoresis of pCDHdp and pCDHsp plasmid DNA extracted from *E. coli*. Lane M, DNA ladder; Lane 1, pCDHdp plasmid DNA; Lane 2, pCDHdp plasmid DNA digested with XbaI and EcoRI enzymes; Lane 3, pCDHsp plasmid DNA; Lane 4, pCDHsp plasmid DNA digested with XbaI and EcoRI enzymes.

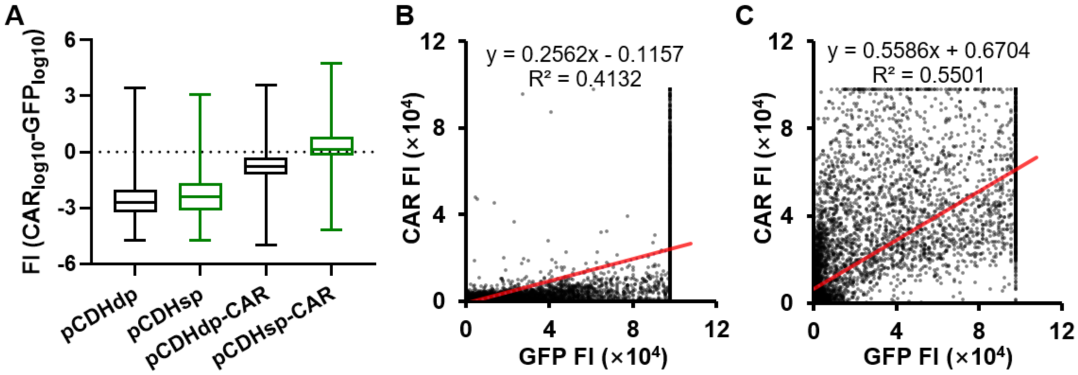


**Fig. S2**. Relationship between the CAR expression level and the GFP fluorescence level in the HEK-293T cells transfected by pCDH vectors. (A) Distribution of the difference ($CAR_{log10}$ minus $GFP_{log10}$) between the fluorescence intensity of CAR and GFP fluorescence in transfected HEK-293T cells on a log10-log10 scale. (B-C) Relationship between the CAR expression level and the GFP fluorescence level in the HEK-293T cells transfected by two CAR-carrying pCDH vectors. FI, fluorescence intensity.

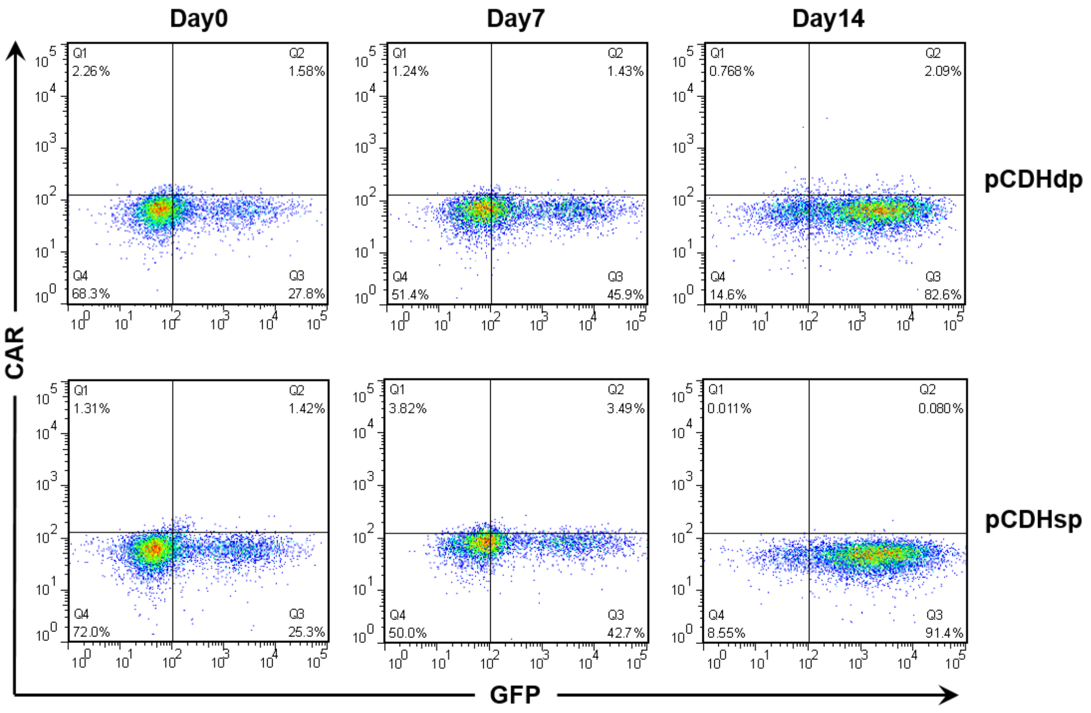


**Fig. S3**. Flow cytometric analysis of GFP and CAR in the NK92 cells transfected by pCDHdp and pCDHsp during puromycin screening.

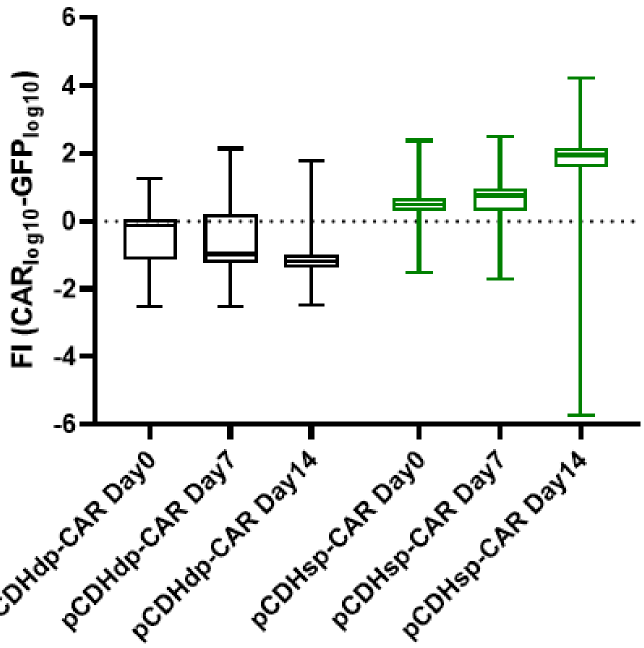


**Fig. S4**. Distribution of the difference (CAR$_{log10}$ minus GFP $_{log10}$) between the fluorescence intensity of CAR and GFP fluorescence in transfected NK cells on a log10-log10 scale during puromycin screening. FI, fluorescence intensity.

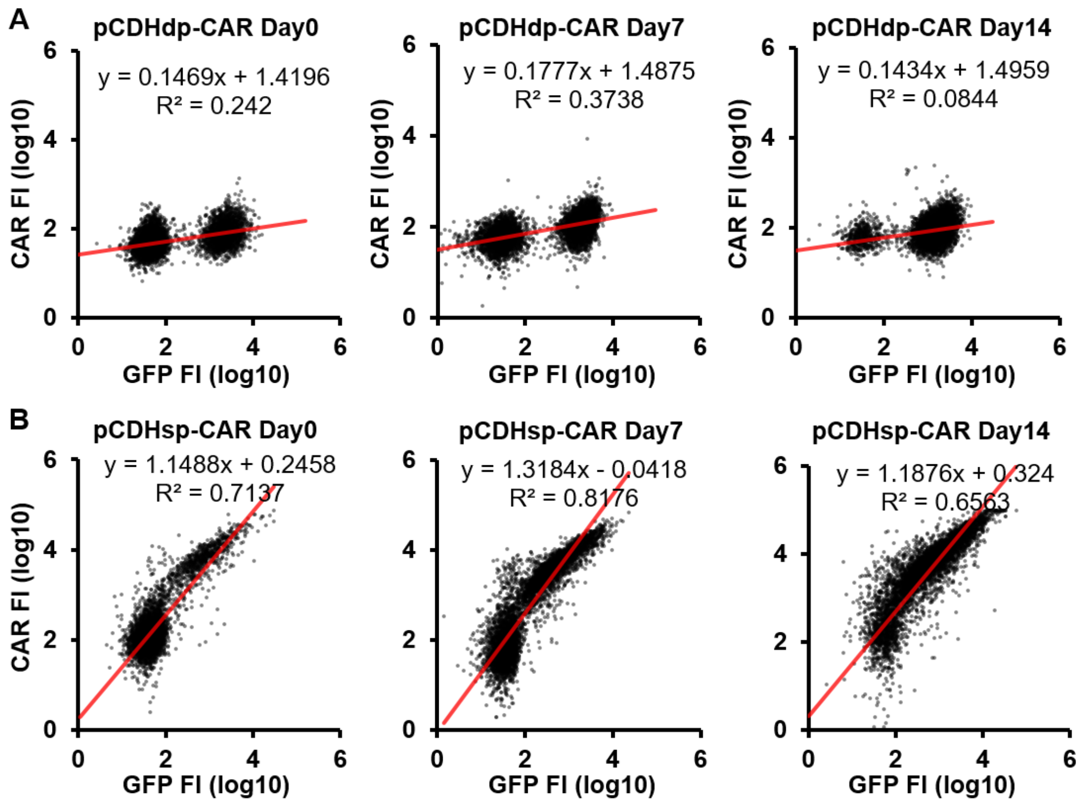


**Fig. S5**. Relationship between the CAR expression level and the GFP fluorescence level on a $\log_{10}$-$\log_{10}$ scale in the NK92 cells transfected by two CAR-carrying pCDH vectors. FI, fluorescence intensity.

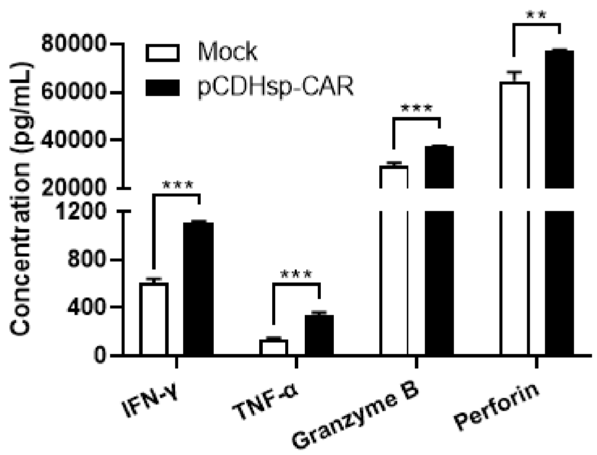


**Fig. S6**. The production of IFN-γ, TNF-α, granzyme B, and perforin were assessed by ELISA in untransduced NK92 cells and pCDHsp-CAR-transduced NK92 cells after stimulation of SGC-7901 cells. Data are shown as mean ± SD. Statistical analysis was done by two-tailed Student's t-test; **$p < 0.01$, ***$p < 0.001$.
